# First detection and molecular identification of the zoonotic *Anaplasma capra* in deer in France

**DOI:** 10.1101/482851

**Authors:** Maggy Jouglin, Barbara Blanc, Nathalie de la Cotte, Katia Ortiz, Laurence Malandrin

## Abstract

Cervids are known to be reservoir of zoonotic tick-transmitted bacteria. The aim of this study was to perform a survey in a wild fauna reserve to characterize *Anaplasma* species carried by captive red deer and swamp deer. Blood from 59 red deer and 7 swamp deer was collected and analyzed over a period of two years. A semi-nested PCR that targets the *23S* rRNA was performed to detect and characterise *Anaplasma* spp. and determine zoonotic species presence. *Anaplasma phagocytophilum* was identified in 14/59 deer (23.7%) but not in swamp deer. Few sequences could not be assigned to any particular species based on the *23S* rRNA sequences. Nested PCR targeting *16S* rRNA, *gltA* and *groEL* genes and sequencing analysis detected a recently reported zoonotic species, *Anaplasma capra* in red deer as well as in swamp deer. This is the first reporting of the tick-borne zoonotic bacterium *A. capra* in France, a species otherwise described only in China and Japan, in goats, sheep, deer and japanese serows. Even if this bacterium may have been introduced in the Park with infected imported animals, its local epidemiological cycle through tick transmission seems possible as locally born deer were found infected. Diagnostic methods, especially molecular ones, should take into account the potential infection of animals and humans with this species.

## Introduction

Bacteria of the genus *Anaplasma* are obligate intracellular parasites that replicate within the vacuoles of diverse eukaryotic cells (monocytes, granulocytes, erythrocytes, endothelial cells). These bacteria are mainly transmitted by ixodid ticks and multiply both in the invertebrate and vertebrate hosts [1]. The genus *Anaplasma* includes 6 recognized species (*A. phagocytophilum, A. bovis, A. centrale, A. marginale, A. ovis* and *A. platys*) responsible for anaplasmosis worldwide on a large range of wild and domesticated vertebrates [1]. One species, *A. phagocytophilum*, described in 1994 in the USA as the agent of human granulocytic anaplasmosis (HGA), is now increasingly detected worldwide [1]. In 2015, a second zoonotic and new species, proposed as *A. capra*, was described in humans in China [2]. On a population of 477 patients with tick-bite history, six percent (28 patients) were found infected with *A. capra* with non-specific febrile manifestations. Five of them were hospitalized due to severe symptoms. General clinical features in human patients have included febrile manifestations (fever, headache, malaise) as well as eschar, lymphadenopathy and gastrointestinal symptoms [2].

Both *A. phagocytophilum* and *A. capra* infect diverse domestic (sheep and goats) and wild ruminants (deer) species, which are considered as reservoirs. In a survey of tick-borne diseases in captive and protected deer in France, we investigated the presence of *Anaplasma* species infecting captive red deer (*Cervus elaphus*) and swamp deer (*Rucervus duvaucelii*). In the “Réserve de la Haute Touche”, endangered species such as the swamp deer (CITES appendix I) are preserved. This reserve is surrounded by a large forested and humid area, a biotope favorable to Ixodid ticks, vectors of *A. phagocytophilum*.

## Methods

### Animal sampling

In 2015, a molecular survey of *Anaplasma* spp. infecting deer was started in the “Réserve de la Haute Touche” Zoological Park, Indre, France (National Museum of Natural History). Blood samples from 59 red deer and 7 swamp deer were collected between 2015 and 2017. They were used for molecular detection and characterization of *Anaplasma* spp.. Blood was sampled at the jugular vein at the occasion of animal care (treatments, vaccinations) or transfers (authorization 36-145-002).

### Molecular detection and characterization of *Anaplasma* spp

Genomic DNA was extracted from blood according to previously described protocols [3]. We detected *Anaplasmatacae* by semi-nested PCR based on the *23S* rRNA gene [4] and determined the species by sequencing PCR positive amplicons. A new detected *Anaplasma* species was further characterized using nested PCR and sequencing of the *16S* rRNA, *gltA* and *groEL* genes (table 1). Bidirectionnal sequencing was performed to ensure sequences, that were further analyzed by the BLASTN (http://www.ncbi.nlm.nih.gov/BLAST/) and CLUSTAWL (https://www.ebi.ac.uk/Tools/msa/clustalo/) programs.

**Table 1.**
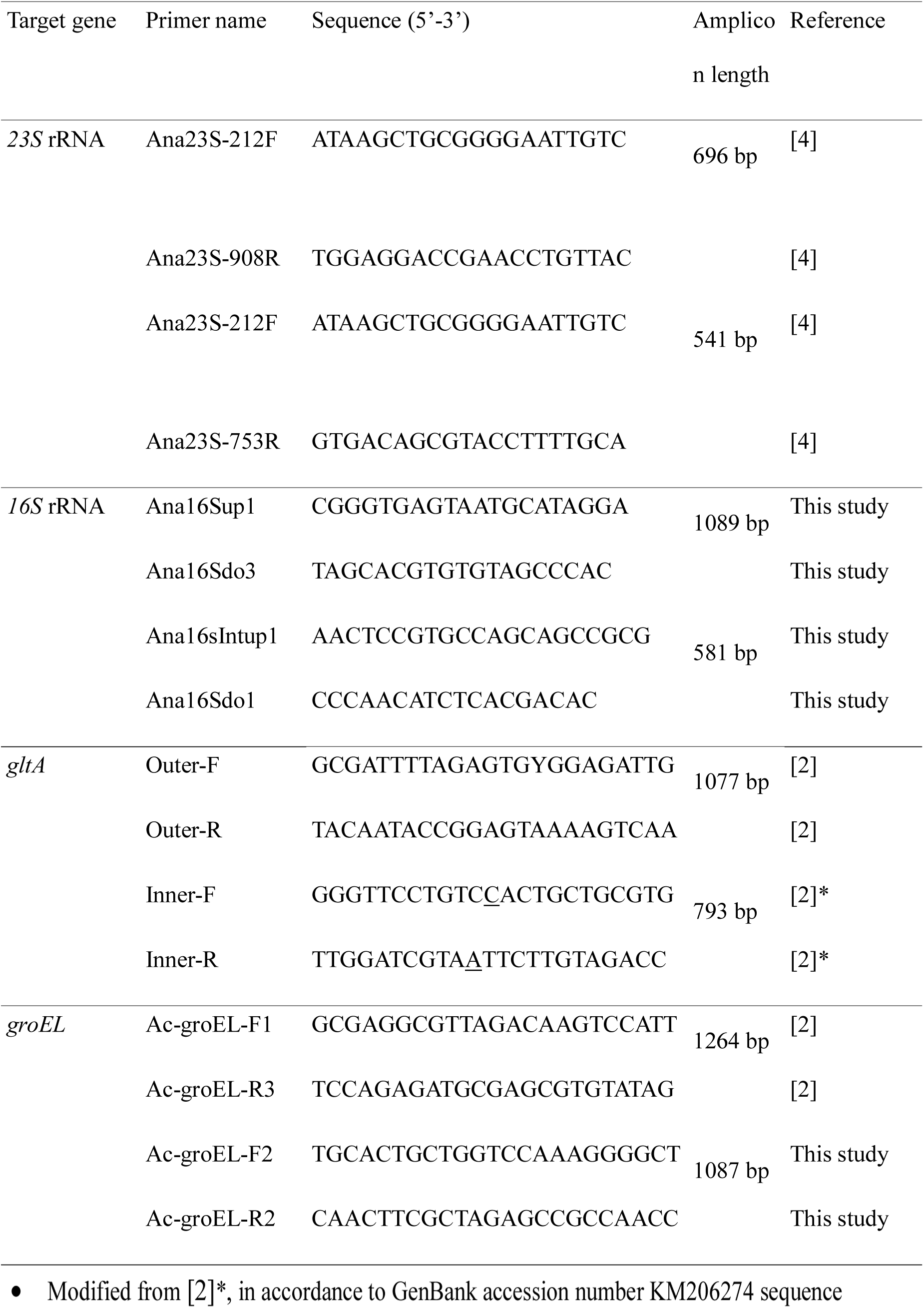
Nucleotide sequence of primers used in the study

## Results

### Detection of Anaplasma spp. in deer blood

Of the 66 heparine blood samples from red deer and swamp deer, *23S* rRNA amplicons of the right size were obtained for 28 samples. A BLASTn search of *23S* rRNA sequences identified *A. phagocytophilum* in 14 red deer (4/21 in 2015, 7/23 in 2016 and 3/15 in 2017) (prevalence of 23.7%) but not in swamp deer. Sequences (lengths between 430-476 bp) were more than 99.5% identical (maximum two mismatches) to *A. phagocytophilum* strain HZ (GenBank accession number NR_076399). Sequences from eleven amplicons (367 to 476 bp) were identical, with 99.8% identities with *Ralstonia pickettii* (GenBank accession number CP001644). Three identical sequences from two red deer and one swamp deer (3/66 - infection rate 4.5%) gave the highest identities with “*Candidatus Anaplasma mediterraneum”* sequence (KY498330), described as a potentially new *Anaplasma* species infecting sheep in Corsica [5] (table 2). Other genetic markers often used to identify *Anaplasma* species were then tested to further characterize this *Anaplasma* species never described in deer.

**Table 2.** Sequence lengths and identity rates with *Anaplasma spp.* sequences

| Gene | Sequence lengths | GenBank accession numbers | Reference organisms and sequences | Identity rate |
| --- | --- | --- | --- | --- |
| 23S rRNA | CEL15 - 367 bp<br>CEL17 - 506 bp<br>DVC 17 - 515 bp | MH084724<br>MH084723 | *Cand. A. mediterraneum* KY498330<br>*A. ovis* KM021411<br>*A. centrale* NR-076686<br>*A. marginale* KY498332<br>*A. platys* KM021412<br>*A. phagocytophilum* KM021418 | **99.6 %**<br>95.0%<br>94.6%<br>93.8%<br>91.6%<br>90.8% |
| 16S rRNA | CEL17 - 531 bp<br>DVC 17 - 518 bp | MH084721<br>MH084722 | *A. capra* KM206273 - MF066917<br>*A. marginale* AF414874<br>*A. centrale* AF318944<br>*A. ovis* AJ633049<br>*A. phagocytophilum* NR-044762<br>*A. platys* AY077619 | **99.6% - 99.8%**<br>98.7%<br>98.5%<br>98.3%<br>96.4%<br>96.2% |
| groEL | CEL15 - 559 bp<br>CEL17 - 1008 bp<br>DVC 17 - 1087 bp | MH084718<br>MH084717 | *A. capra* KM206275 - AB454078<br>*A. marginale* AF414864<br>*A. centrale* EF520691<br>*A. ovis* AF441131<br>*A. platys* AY044161<br>*A. phagocytophilum* JF494833 | **91.4% - 97.7%**<br>83.2%<br>83.2%<br>83.2%<br>77.5%<br>76.2% |
| gltA | CEL15 - 729 bp<br>CEL17 - 707 bp<br>DVC 17 - 725 bp | MH084720<br>MH084719 | *A. capra* KM206274 - MG940872<br>*A. marginale* AF304139<br>*A. ovis* PKOE01000003<br>*A. centrale* CP001759<br>*A. phagocytophilum* AY464132<br>*A. platys* EU516387 | **87.9% - 98%**<br>74.6%<br>74.2%<br>73.3%<br>65.4%<br>61.4% |

CEL : *Cervus elaphus*
DVC : *Rucervus duvaucelii*

### Further molecular characterization of the new Anaplasma spp. from deer in France

Two *16S* rRNA identical sequences were obtained from red and swamp deer blood samples. They had similarities of more than 99.6% with numerous *16S* rRNA *Anaplasma capra* sequences deposited in GenBank. These sequences were obtained from sheep, goat, human blood and ticks from China, as well as from cattle, sika deer, Japanese serows and ticks from Japan. Sequence similarities with other known *Anaplasma 16S* rRNA sequences were lower than 99% (table 2).

As *Anaplasma capra groEL* and *gltA* sequences were also deposited in GenBank, we amplified and sequenced these genes from our deer blood samples to better characterize this new *Anaplasma*. The three *groEL Anaplasma* sequences from deer were identical and identity rates ranged from 91.4 to 97.7% with the *A. capra groEL* sequences from China and Japan available in GenBank (table 2). The similarities with *groEL* sequences from other related *Anaplasma* species (*A. centrale, A. marginale, A. platys, A. phagocytophilum* and *A. ovis*) fell under 84%. The three *gltA Anaplasma* sequences from deer differed by one nucleotide. The identity rates of the longest sequence (729 bp) ranged from 87.9 to 98% with *A. capra gltA* sequences from Japan and China. They were lower than 75% (61.4 to 74.6%) with *gltA* sequences from other *Anaplasma* species (table 2). All these data confirmed the identity of the *Anaplasma* from French deer as belonging to the *A. capra* species.

Partial sequences of the *16S* rRNA, *23S* rRNA, *groEL* and *gltA* from *A. capra* identified from the swamp deer and one red deer were deposited in GenBank (accession numbers MH084717- MH084724 with details in table 2).

### Persistence of A. capra

The persistence of deer infection by *A. capra* was analyzed by sampling blood from one of the two infected red deer four months after the initial detection of this unexpected bacterial species. We detected *A. capra*, with *23S* rRNA, *16S* rRNA, *groEL* and *gltA* sequences 100% identical to the first identified *A. capra*, with the same distinct nucleotide in the *gltA* sequence as characterized 4 months earlier.

## Discussion

For about 40% of the positive conventional Anaplasmataceae spp. specific semi-nested PCRs, sequences indicated amplification of *Ralstonia pickettii 23S* rRNA, highlighting a lack of specificity of the semi-nested PCR used. As some of our negative controls were also positive, we decided to sequence them and sequences corresponding also to *Ralstonia picketii* were found. As this bacteria is a frequent contaminant of all kind of solutions, including ultrapure water [6], our results most probably correspond to *R. pickettii* contamination of the solutions we used for extraction or PCR.

In this survey, we detected two *Anaplasma* species infecting deer. *A. phagocytophilum* was detected with a rather moderate prevalence (23.7%) in red deer only. Prevalence of *A. phagocytophilum* infection in wild red deer in Europe is highly variable, from 1.5% in Austria, 10.9% in Portugal, 40-75% in Italy, 80.8% in Spain, to 97.9 to 100% in central Europe (respectively in Slovakia and Hungary) [7–13]. Captive deer are probably less prone to tick bites compared to wild ones, due to grazing areas management. This result indicates anyway the contact of red deer with ticks and the transmission of *A. phagocytophilum* in the Reserve. Swamp deer were found uninfected with *A. phagocytophilum*, a result which could be attributed to the low number of animals analyzed in the case of a low infection rate (7). There are no data about tick-transmitted pathogens for this endangered species, so the susceptibility of swamp deer to *A. phagocytophilum* is unkown. A recently described *Anaplasma* species, *A. capra* was detected and identified in both deer species, with a much lower infection rate (4.5%). *A. capra* has already been detected in various ruminant hosts (sheep, goats, cattle, sika deer, Japanese serows) but its localization was up to now geographically restricted to China and Japan [14–17]. Human infection by this newly-described species has been reported in northeast China, leading to patients hospitalisation [2]. The detection and characterization of *A. capra* based on several molecular markers in our study represents the first evidence of this potentially new zoonotic species in Europe (France) in two new hosts, red deer and swamp deer. Species assignation to *A. capra* was based on *16S* rRNA homologies higher than 99% [18].

The *23S* rRNA sequence from *A. capra* described in our study blasted with an unknown *Anaplasma* species proposed as “*Candidatus Anaplasma mediterraneum*” from sheep in Corsica (France)[5]. Whether “*Candidatus Anaplasma mediterraneum*” corresponds in fact to *A. capra* could not be determined, as the only other marker used in this study is *rpoB*, whereas we as most authors used a combination of *16S* rRNA, *groEL, gltA* and *msp4* sequences to identify and characterize *Anaplasma capra* [2, 15–17, 19–21].

We have detected *A. capra* in three different deer since 2015. The first infected and detected red deer was a male originating from France (Theix) while the two others (red deer and swamp deer) detected in 2017 and 2018 were both born inside the Park. Acquisition of a locally transmitted *A. capra* is therefore probable for these two deer, even if *A. capra* may have been originally introduced into the Park from an external source. The epidemiological cycle of *A. capra* seems therefore to be completed locally. The low prevalence of infected deer in the Park might be due to the introduction being recent. Ticks are the main vectors for *Anaplasma* species even if other transmission routes have been described for some species (blood-sucking flies and transplacental transmissions) [1]. Although *A. capra* has been detected in several tick species, *Ixodes persulcatus* [2], *Rhipicephalus microplus* [19], *Haemaphysalis longicornis* [15,20] and *Haemaphysalis qinghaiensis* [21], vectorial competence has not been proven yet. As most of these tick species are not present in France, another tick species may be responsible for *A. capra* transmission in France. The “Réserve de la Haute Touche” is located in a forested preserved area suitable for ticks and ticks are commonly found feeding on the animals as well as questing on the vegetation (not shown). Vector identification and vectorial competence remain to be elucidated.

In this study, we demonstrated the presence in France of the new species *A. capra* on two new hosts. New studies are required, to examine its zoonotic ability, as non-zoonotic genetic variants may exist as described in the case of *A. phagocytophilum* [1,3]. Diagnostic methods, especially molecular ones, should take into account the potential infection of animals and humans with this species, as molecular tools are often designed to specifically detect *A. phagocytophilum.* To improve our knowledge on the epidemiological cycle of this bacterium in France, the vector tick species should be identified, in order to evaluate the risks of transmission to humans. Deer could therefore be considered as a potential reservoir for *A. capra*.

## Data Availability Statement

Gene sequences are available in GenBank under accession numbers MH084717-MH084724.

## Acknowledgements

Special thanks go to Alice Brunet, Héloïse Duchêne, Emmanuel Maréchal and Christophe Jubert for handling the deer and for their help in this survey as well as to the entire staff of the Zoological Park of “La Haute Touche” for their care of the animals. Roland Simon is also acknowledged for making this study possible.

## Author Contributions

**Conceptualization:** Laurence Malandrin, Katia Ortiz

**Formal Analysis:** Maggy Jouglin, Laurence Malandrin

**Funding Acquisition:** Maggy Jouglin, Nathalie de la Cotte, Laurence Malandrin

**Investigation:** Barbara Blanc, Maggy Jouglin, Nathalie de la Cotte

**Project Administration:** Barbara Blanc, Katia Ortiz, Laurence Malandrin

**Resources:** Katia Ortiz, Barbara Blanc, Nathalie de la Cotte

**Supervision:** Katia Ortiz, Laurence Malandrin

**Visualization:** Katia Ortiz, Laurence Malandrin

**Writing – Original Draft Preparation:** Maggy Jouglin, Laurence Malandrin

**Writing – Review & Editing:** Barbara Blanc, Maggy Jouglin, Katia Ortiz, Nathalie de la Cotte, Laurence Malandrin

